# Whole-genome resequencing identified loci underwent divergent selection and improved local adaptability in groundnut (*Arachis hypogaea*)

**DOI:** 10.64898/2026.07.13.738128

**Authors:** Muhammad Jahanzaib, Kunhui He, Sana-ur Rehman, Ihsanullah Khan, Haris Khurshid, Muhammad Jawad Umer, Sunil Gangurde, Awais Rasheed, Huihui Li

**Affiliations:** Department of Plant Sciences, Quaid-i-Azam University, Islamabad 45320, Pakistan; State Key Laboratory of Crop Gene Resources and Breeding, Institute of Crop Sciences, Chinese Academy of Agricultural Sciences (CAAS), CIMMYT-China Office, Beijing, 100081 China; Nanfan Research Institute, CAAS, Sanya, Hainan, 572024 China; Oilseed Research Program, Crop Sciences Institute, National Agricultural Research Center, Islamabad, Pakistan; Crops Research Institute, Guangdong Academy of Agricultural Sciences, Guangdong Provincial Key Laboratory of Crop Genetic Improvement, South China Peanut Sub-Center of National Center of Oilseed Crops Improvement, Guangzhou, Guangdong Province, 510640, China; College of Agronomy, Shandong Agricultural University, Tai’an 271018, China

**Author notes:** Authors contributed equally (Muhammad Jahanzaib, Kunhui He).

**Keywords:** Groundnut, Whole-genome resequencing, Genome-wide association studies, Selective sweeps, SNP markers

## Abstract

There is an urgent need to expand groundnut genomics knowledge base to improve yield and adaptability in the target environments. Whole-genome sequencing can discover selective sweeps in the genomic regions distinguishing adaptive from non-adaptive germplasm within target environments. When combined with genome-wide association studies (GWAS), this approach can reveal genes underpinning local adaptability and yield advantage. The objective of this study was to establish a genome-wide quantitative framework form identifying genomic regions under selection, with a particular focus on narrowing down regions associated with important yield and adaptive traits. Moreover, validation of elite haplotype distributions in an independent fully sequenced groundnut panel. A panel of 197 groundnut accessions was subjected to whole-genome sequencing and phenotypic evaluation to dissect the collection into adaptive and non-adaptive subsets to uncover the genomic regions under selection. This was then combined with GWAS to uncover genetic variants governing agronomic traits associated with yield and adaptability. The stringent single and multi-trait analysis identified 60 loci for 12 agronomic traits, of which seven loci controlled multiple yield related traits and were pleiotropic. Within our diversity panel, 48 genomic regions showed signs of selection. Among these selective sweeps, six positively selected loci were co-localized with trait associated loci. A large genomic region on chr2 spanning ∼78 Mb was under divergent selection and harbored genes underpinning yield and 20-pod length. A F-box transcription factor, *Arahy.37HYKA,* on chr9, and an alanine transferase protein gene, *Arahy.E9MTVL,* on chr12 carried peak SNPs associated with yield and related traits. We further cross-validated our results in another groundnut355 panel, where the corresponding genes within LD blocks showed significant effects on HKW, pod length and pod weight. The genomic resources developed here provide a high-resolution variation map to delineate the genes underpinning yield and developmental traits in groundnut, improved our understanding of the genetic basis of important agronomic traits, and provide a valuable resource for further functional genomics studies and groundnut improvement programs.

## Introduction

Groundnut commonly referred to as peanut (*Arachis hypogaea* L.) is an important oil and food legume crop worldwide (Cason et al., 2023). Groundnut is not only important source for vegetable oil but is also a major cash crop for elevating farmers income. The globally cultivated area of groundnut was registered 32.7 m ha with 53.9 m tons annual production, out of which 90% of both groundnut areas and global production is from developing countries (FAOSTAT, 2024). Groundnuts offer a rich source of healthy mono-, and poly-unsaturated fatty acids that plays vital role in cholesterol regulation as well as supporting cellular health (Shah et al. 2023). The demand of groundnut and groundnut based products is high due to its high oleic and linoleic acid, fiber, folic acid, and certainly digested protein, as well as the presence of resveratrol (Raza et al. 2024). Groundnuts originated from tropical and subtropical regions of South America, introduced to Spain in 16^th^ century and were widely cultivated in many European countries until 1840. Groundnut production remains low in several parts of the world, and several reasons of low productivity are biotic and abiotic stresses, ineffective seed distribution techniques, lack of scientific knowledge, economic growth, low-input utilization, and political and social facilities (Varshney et al. 2017).

Population genomics facilitates uncovering the numerous loci involved in the evolutionary processes and breeding progress, thereby unfold the mechanisms of inbreeding and outbreeding depressions, population demographic history and genetic basis of local adaptation (Bhat et al 2021). Genome-wide association studies (GWAS) have been used in the past for linking molecular markers and QTLs associated with significant attributes in groundnut (Otyama et al. 2022; Pandey et al. 2014). In this context, the availability of a good quality reference genome, and genome resequencing at the population level are important strategies to underpin the loci associated with local adaptability. The international groundnut genome initiative and other groups have released reference genomes of cultivated groundnut (Bertioli et al. 2019; Chen et al. 2019; Zhuang et al. 2019), and their progenitors (Bertioli et al. 2016; Chen et al. 2016b). Likewise, the whole-genome resequencing of large groundnut populations has resulted in identification of loci underpinning agronomic traits evolutionary history (Lu et al. 2024; Zheng et al. 2024). The development of whole-genome re-sequencing knowledge has made to secure high-throughput SNP markers progressively more feasible, providing robust support for inspecting the molecular-level genetic basis of quantitative traits (Zhang et al., 2023).

Previously, several GWAS have been conducted for consolidate mapping of very complex traits in populations like pod weights (Gangurde et al. 2020), yield (Guo et al. 2024), growth habit (Li et al. 2022), resistance to leaf spot (Oteng-Frimpong et al. 2023), and fatty acid composition (Otyama et al. 2022) are some of significance of GWAS in groundnuts. A major limitation in these studies is the use of low-density SNP platforms like reduced representation sequencing or SNP arrays. The use of whole-genome resequencing could solve this issue. Modern crop breeding activities select favourable alleles in new cultivars, and these signatures of selection can be detected by a cross-population comparison approach like EigenGWAS (Chen et al. 2016a). Several lines of evidence have shown that that genomic regions that exhibit selection signatures are also enriched for genes associated with biologically important traits (Xie et al. 2015). Therefore, detection of selection signatures is emerging as an additional approach to identify and validate novel gene-trait associations in the modern or adapted germplasm in wheat (Afzal et al. 2019), barley (Sharma et al. 2021), and maize (Li et al. 2019; Li et al. 2021).

This is the first study on the whole-genome resequencing of groundnut germplasm adapted in Pakistan and provide deep insight into the genomic regions underpinning agronomic traits and local adaptability.

## Materials and Methods

### Plant material and field trials

A diversity panel comprised of 197 accessions were used in this study (Supplementary Table S1). These 7 cultivars and 128 accessions from four local breeding programs, and 68 exotic accessions from USDA, and ICRISAT. The local accessions were collected from Oilseed Research Program (NARC), Plant Genetic Resources Institute (NARC, Pakistan), Groundnut Research Station (Attock, Pakistan) and Barani Agriculture Research Institute (Chakwal, Pakistan).

The research was conducted over two consecutive years (2021 and 2022). The experiment was performed at the Oilseeds Research Program at NARC Islamabad (33.6701° N, 73.1261° E) under optimal field conditions recommended for groundnut cultivation. The seed were planted in Alfa Lattice Design with three replications. The sowing dates were April 7 and May 5 during the Kharif seasons of 2021 and 2022, respectively. The experimental plot size was consisted of three rows of 4 meters length with row-to-row distance of 45 cm, and plant-to-plant distance was maintained at 10 cm for proper plant growth. The standard agronomic practices were followed. The data was recorded on 12 morphological traits related to architecture including plant height (cm), plant width (cm), number of branches, stem girth (mm), leaflet length (unit), leaflet width (unit), and the remaining six were yield related traits including 20-pod length (cm), shelling percentage (%), 100-kernel weight (g), kernel length (cm), kernel width (cm), and yield (kg/ha).

### Phenotypic data analysis

The analysis of variance (ANOVA) was used to test the statistical significance of different sources of variation for each of the twelve traits. In the ANOVA model, phenotypic effect was partitioned into overall mean, year (environment) effect, replication (i.e. block) within year, genotypic effect, genotype by environment effect, and random error effect. Let *y*_ijk_ be the observed value of a trait of interest for the *i*^th^ accession in the *k*^th^ replication under the *j*^th^ environment (equivalent to year in this study). The linear model used in ANOVA is therefore,

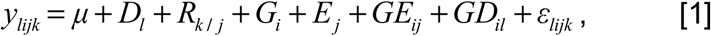

where *i* = 1, 2, …, *n* (*n* = 192), *j* = 1, 2, …, *e* (*e* = 4 with two locations and two years), *k* = 1, 2, …, *r* (*r* = 2), is overall mean of the whole population, *R*_k/j_ is the *k*^th^ replication effect in the *j*^th^ environment, *G*_i_ is genotypic effect of the *i*^th^ accession, *E*_j_ is environmental effect of the *j*^th^ environment, *GE*_ij_ is interaction effect between the *i*^th^ accession and the *j*^th^ environment, *GD*_il_ is interaction effect between the *i*^th^ accession and the *l*^th^ treatment, and ε_lijk_ is random error effect which was assumed to be normally distributed with a mean of zero, and variance 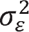. The ANOVA described above was implemented with the GLM procedure in SAS software (SAS Institute, Cary, NC, 2007).

The BLUP of genotypic value for each accession under each water treatment was used as the phenotype for all subsequent comparisons. BLUPs were calculated as follows: the observed value of trait was defined as *y*_ijk_ for the *i*^th^ accession in the kth replication in the *j*^th^ environment (equivalent to location and year in this study). The mixed model used for BLUP was therefore,

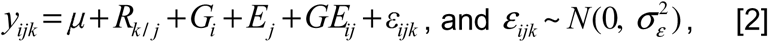

where *i* = 1, 2, …, *n* (*n* = 203), *j* = 1, 2, …, *e* (*e* = 4 with two locations and two years), *k* = 1, 2, …, *r* (*r* = 2), *R_k/j_*, *G_i_*, *E_j_*, and *GE_ij_* were the same as the description above. Except for, all the effects were viewed as random effects following the normal distributions 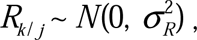 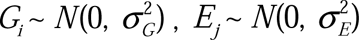 and 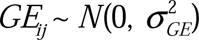, where 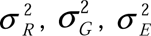, and 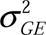 were the variances explained by replication, genotype, environment, and genotype by environment interaction, respectively. The BLUPs were calculated with the MIXED procedure in SAS software (SAS Institute, Cary, NC, 2007).

The Pearson’s coefficient of correlation was calculated using corrplot program implemented in R v. 4.3.1.

### DNA isolation and sequencing

The genomic DNA was extracted from 3-5 cm long seedling leaves grown in peatmoss filled plastic trays. The DNA was extracted using Plant Genomic DNA Extraction Kit (TIANGEN, Beijing, China) according to the instructions of the manufacturer (http://www.tiangen.com/en/?productShow/t1/4/id/10.html). After extraction, DNA was quantified on 1% agarose gel, and a quantity of 100 ng/μl was maintained for subsequent SNP analysis.

### Whole-genome resequencing and variant calling

The paired-end sequencing libraries were constructed with inserts of approximately 300 bp and sequenced with Illumina HiSeq 6000 platform with PE150. The raw data generated were cut with average coverage of 5x per for further analysis. The reads were trimmed and subjected to quality-control for further alignment to reference genome. The subsequent high-quality reads were aligned to the genome of cultivated *A. hypogaea* cv. Trifrunner version 2 (https://data.legumeinfo.org/Arachis/hypogaea/genomes/Tifrunner.gnm2.J5K5/) using burrows-wheeler alignment tool (BWA) V0.7.17 software with default parameters. The SNPs and InDels were called using Genome Analysis Tool Kit (GATK) pipeline with the haplotype caller pipeline (Poplin et al. 2017). Briefly, the SAMTOOLS V1.7 software to convert the sequence alignment/map (SAM) file generated in the first step into binary alignment map (BAM) format and perform sorting, and genome analysis toolkit (GATK) V4.2.1.0 software was used to analyze and classify the SNP. To generate the integrated VCF data, we merged SNP marker using a custom R script, according to overlapped chromosome and physical position. Moreover, SNP data were further filtered using VCFtools V0.1.16 (Danecek et al. 2011). The SNPs and InDels in the above steps were again filtered for further analysis. All the biallelic SNPs and InDels with missing rate >0.05, minor allele frequency <0.05, and number of heterozygous genotypes >20 were filtered using bcftools (v 1.10.2) and a final vcf file was generated for subsequent analyses.

### Linkage disequilibrium and population structure

Population structure and cryptic relatedness of the 197 groundnut accessions were inferred using admixture analysis, principal component analysis (PCA), and a neighbor-joining phylogenetic tree. The population structure was calculated using a Bayesian approach-based program implemented in fastSTRUCTURE with the number of population clusters (K) set from 2 to 20 (Raj et al. 2014). The optimal K (5) was determined using the chooseK.py script in fastSTRUCTURE. The highest estimated membership coefficient criterion was employed to assign unknown accessions to cluster. PCA based on the same dataset was performed at the individual level using the SNPRelate R package (Zheng et al. 2012). The top two principal components were used to visualize the genetic relationships among groundnut accessions. The PCA biplot was generated using the ggplot2 package in R with pre-defined subgroups. In addition, TASSEL V5 software was used to construct a neighbor-joining tree according to the distance matrix. The tree was visualized using the “iroki” online tool (https://www.iroki.net/).

The patterns of genome-wide linkage disequilibrium (LD) among SNPs were studied as an indicator of further population structure to test selection pressure. The degree of LD between each pair of SNPs was evaluated as squared allele frequency correlation (*r^2^*), estimated by PopLDdecay with default parameters except for -MaxDist 600 (https://github.com/BGI-shenzhen/PopLDdecay). The LD decay was calculated for each subpopulation based on the *r^2^*for each pair of SNPs using a custom R script. The *r^2^* estimates were transformed using a nonlinear square root transformation according to the Hill and Weir model (Hill and Weir 1988). The average genome-wide LD decay plot was visualized using R software (R Core Team 2016).

### Selective sweep analysis by XPCLR

To detect the loci that may have undergone selection in two sub-populations, XP-CLR along with population fixation statistics, *F*_ST_, using vcftools was used. A 100-kb sliding window and a 10-kb step size was used for XP-CLR analysis. To ensure comparability of the composite likelihood score in each window, we fixed the number of SNPs assayed in each window to five with the setting ‘--maxsnps 5 --minsnps 5 (Chen et al. 2010). Meanwhile, to keep the used genomic windows consistent in the XP-CLR analysis, the weighted *F*_ST_ values were estimated in each window that required at least five SNPs with the setting ‘--fst-window-size 100,000 --fst-window-step 10,000. Pairwise differentiation between populations (*F*_ST_) was calculated using the “hierfstat” package of R (Goudet 2005). Evidence for selection across the genome we merged the adjacent windows with top 10% values into a single window, and the top 0.5% outliers were determined to represent putative selection signals. In addition, adjacent sweeps separated by a physical distance of < 100 kb were merged into a single selected locus.

### GWAS analysis

The GWAS was performed for 12 quantitative traits on 197 groundnut accessions using 707138 genome-wide SNP markers. For GWAS, a univariate linear mixed model from the GEMMA software package (Zhou and Stephens 2012). Population structure was rectified using the relatedness matrix and top five principal components. To avoid false-positive associations, a p-value threshold of 1 x 10-5 was chosen to declare marker-trait associations. A multi-trait GWAS was conducted using matrix variate linear mixed model (mvLMM) to identify the pleiotropic loci associated with two group of traits i.e. plant architecture related traits and yield related traits (Furlotte and Eskin 2015).

## Results

### Phenotypic diversity and heritability of key agronomic traits

Comprehensive phenotypic characterization of 12 morphometric traits across 197 groundnut accessions for two consecutive years revealed high phenotypic variation within the panel **(Supplementary Table S2 and Figure 1)**. Most traits showed a normal distribution, except for leaflet width, number of branches and shelling percentage. The plant height averaged 25.5 cm, ranging from 12.7 to 47.2 cm, corresponding to a 3.7-fold variation across years. The yield displayed pronounced diversity, with a mean of 2164 kg ha^-1^ and a range of 453.4-5509 kg ha^-1^, representing a 12-fold variation among accessions. Kernel width, was the least variable trait, varying only 1.4-fold and ranged from 7 to 9.9 mm. The analysis of variance (ANOVA) confirmed highly significant differences among genotypes, years, and their interactions **(Supplementary Table S2)**. Broad-sense heritability varied among traits, ranging from 0.4 for yield (Yld) to 0.88 for kernel width (KW), indicating a substantial genetic distribution to the observed phenotypic variation. Detailed descriptive statistics **(Table)** revealed that yield exhibited the highest coefficient of variation in both years (45.7-49.6%), indicating strong environmental influence, while kernel width showed the lowest CV (12.9-13.0%), demonstrating greater stability across environments. Skewness values confirmed normal distribution for most traits, except number of branches and seeds per pod, which showed positive skewness (1.35-1.38), suggesting improvement potential through selection. Shelling percentage consistently displayed negative skewness (-0.86), indicating favorable clustering toward higher values. Year to year comparisons revealed marginal improvements in yield (2155.5 to 2173.6 kg ha^-1^) and shelling percentage (58.7 to 60.5%), while seed germination showed the most pronounced increase from 4.43 to 5.25.

**Figure 1.**
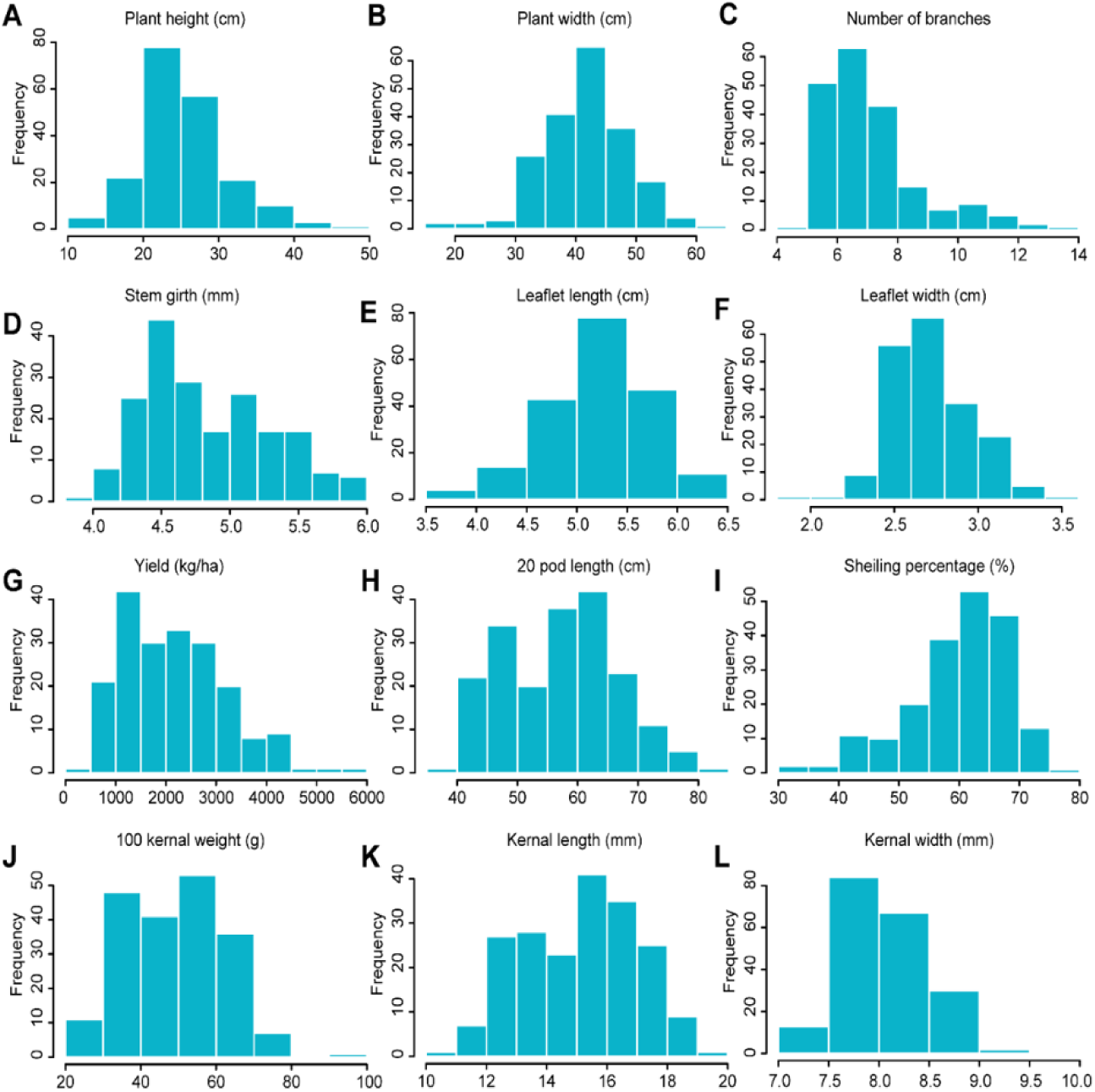
Frequency distribution of 197 groundnut traits based on morphological analysis.

### Correlation analysis of 12 morphological traits

Phenotypic correlation analysis among yield related attributes **(Figure 2)** showed several significant associations. 100-kernel weight (HKW) had a high significant positive correlation with kernel length (0.79***), 20-pod length (0.70***) and yield (0.52***). Conversely, HKW showed significant negative correlations with stem girth (-0.47***) leaflet width (-0.49***), plant height (-0.28***) and leaflet length (-0.21**). Similarly pod yield showed highly significant positive correlations with Kernel width (0.40***), HKW (0.52***), kernel length (0.46***) and 20-pod length (0.49***). Whereas yield showed highly significant negative correlations with stem girth (-0.38***), leaflet width (-0.39***), plant hight (-0.33***) and leaflet length (-0.26***).

**Figure 2.**
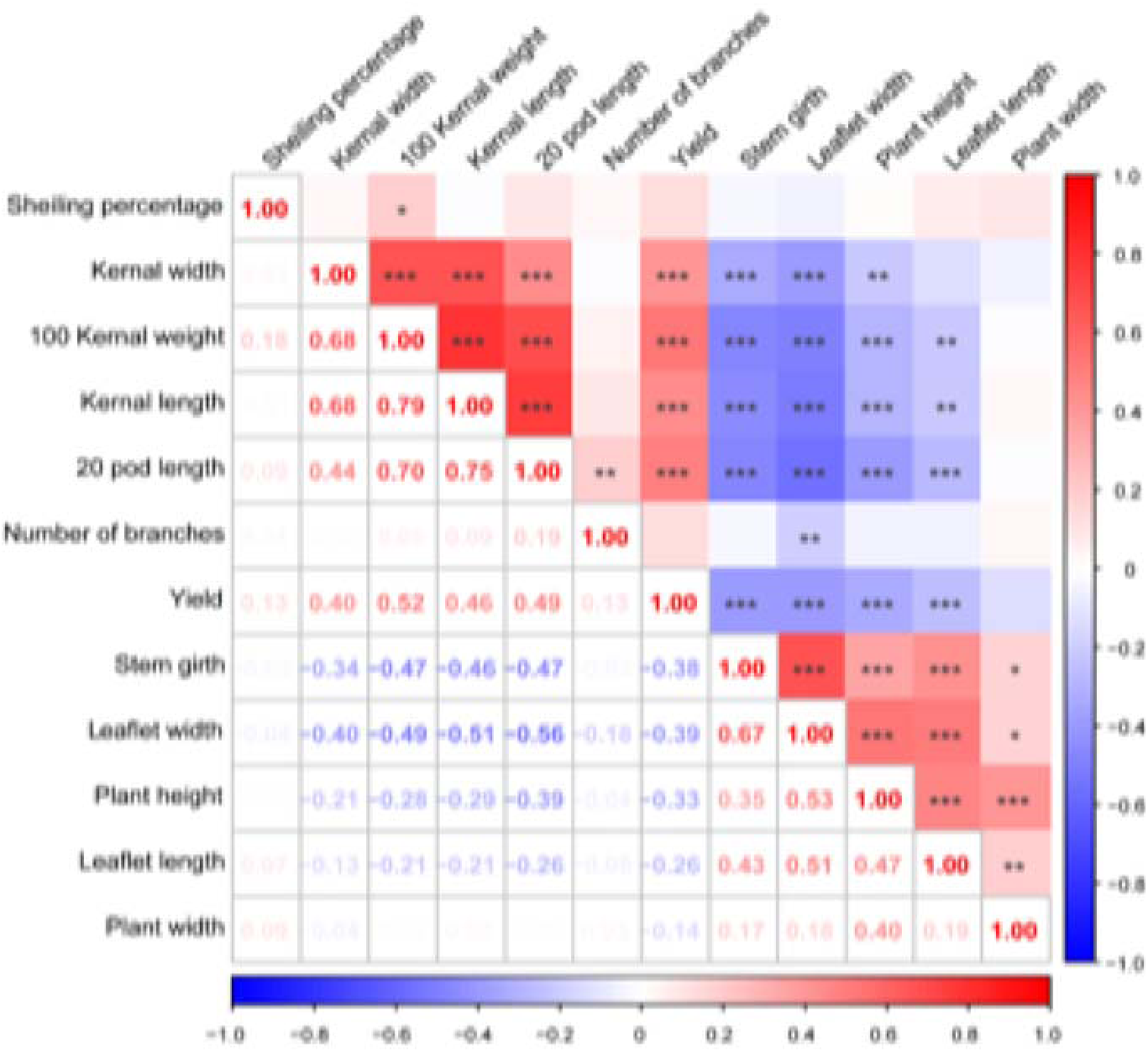
Phenotypic correlation of 12 morphometric traits of 197 groundnut genotypes. Asterisks, *, ** and *** indicates that *P* < 0.05, *P* < 0.01 and *P* < 0.001, respectively.

### SNP distribution in groundnut accessions

To identify candidate genes underpinning important phenotypic traits, a whole-genome genetic variation map was constructed using resequencing data from 197 groundnut accessions. A total of ∼10.4 billion 150-bp paired-end reads were generated. These reads yield a 1.4 Tb of clean data after quality trimming, with an average sequencing depth of 3.1x (ranging from 1.8x-4.9x) **(Supplementary Table S3)**. The filtered and trimmed reads were aligned to the groundnut reference genome (*A. hypogaea* cv. Tifrunner, version 2) identifying ∼1.8 million SNPs and InDels. After filtering for minor allele frequency (MAF > 0.05) and missing date rate (<0.01), 707,138 high quality SNPs were retained for downstream analysis. The descriptive statistic of final genetic variation map revealed variable SNP density across chromosomes **(Figure 3A; Supplementary Table S4).** The highest SNP density per Mb size (350 SNPs) was observed on chr16, whereas the lowest densities were recorded on chr1 (230 SNPs/Mb) and chr15 (237 SNPs/Mb). Across all 20 chromosomes, the total SNP count ranged from 12232 (chr8) and 53213 (chr19) **(Figure 3B)**. The subgenome analysis showed that the B subgenome harbored more SNPs (415,225) compared to the A subgenome (291,913) **(Supplementary Table S3).** Given the total genome size of ∼2.5 Gb, the overall variant density was one SNP every 3580 base pairs. The mean genotype heterozygosity rate was 0.034 with range in between 0 to 0.20 across accessions **(Figure 3C and 3D**). The average MAF across all retained SNPs was 0.227 **(Figure 3E)**. Among all the identified variants, 13661 and 31283 SNPs were intragenic in the exonic and intronic regions, while the remaining SNPs were intergenic **(Supplementary Table S4)**. Among the high impact variants, 356 SNPs were predicted to cause stop_gained mutations, 62 SNPs were classified as stop_lost and 22 SNPs were identified as start_lost mutations, representing putatively deleterious variants that may affect gene function.

**Figure 3.**
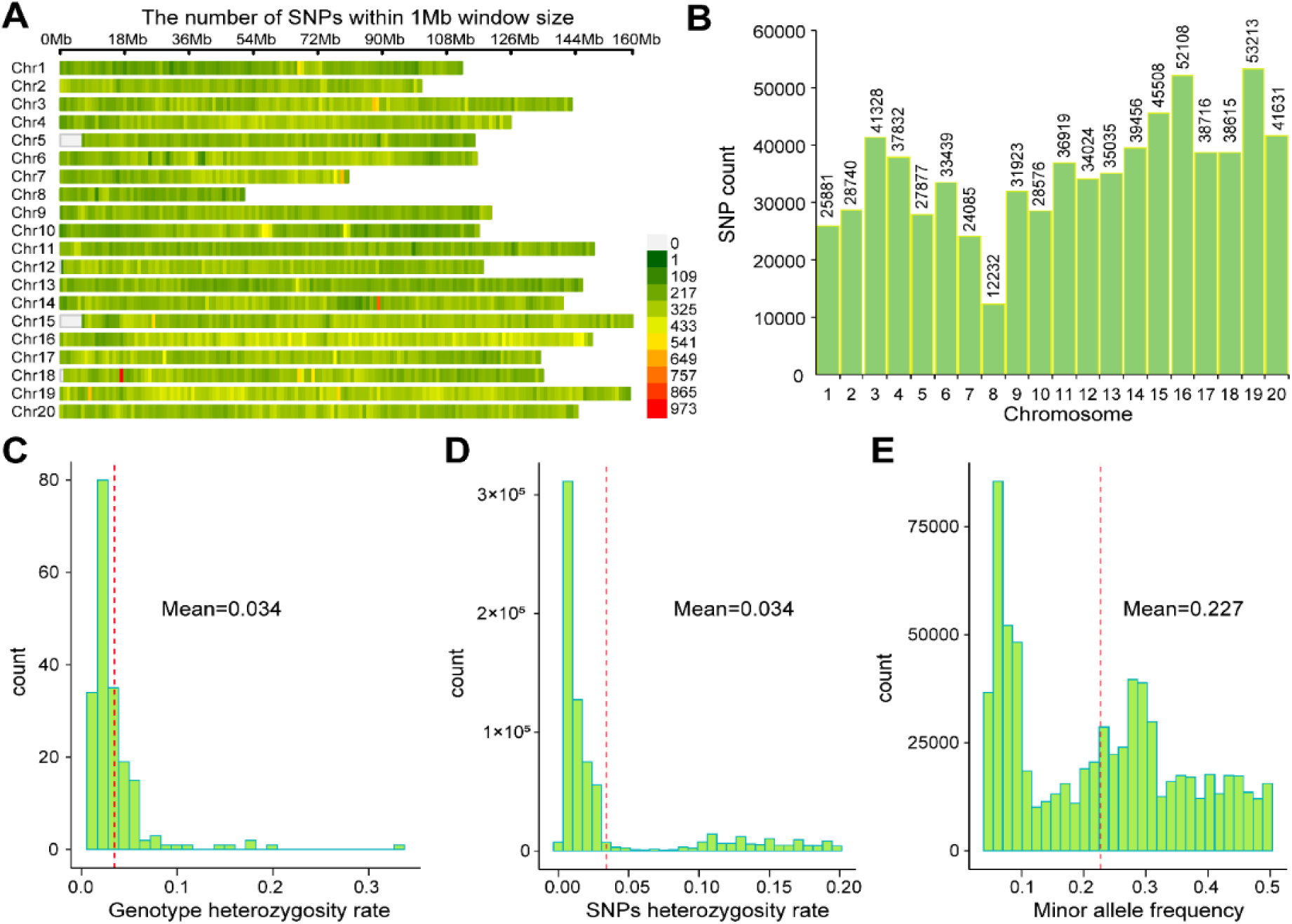
Descriptive statistics of SNPs in groundnut populations. **A** Density distribution of SNPs across the whole genome. **B** The number of SNPs on each chromosome. **C** Population genotypic heterozygous rate. **D** Heterozygosity rate of genome-wide SNPs. **E** Minor allele frequency statistics for genome-wide SNPs.

### Population structure and LD decay in the diversity panel

To elucidate the genetic relationship and population structure among 197 groundnut accessions, the fastSTRUCTURE and principal component analysis (PCA) was used. The population structure analysis revealed two main genetic groups, which were further subdivided into 8 sub-populations (coded: population 1-population 8) **(Supplementary Table S1)**. Group 1 included 143 accessions while Group 2 contained 54 accessions. Most groundnut cultivars bred in Pakistan were included in Group 1, however 26 accessions, including advanced breeding lines developed at NARC, Pakistan were included in Group 2. Notably, most of the accessions from Uruguay, Paraguay and Brazil were included in Group 2. Among the 8 sub populations, population 1 contained the highest number of accessions (n=57), whereas population 3 contained the lowest (n=6) **(Figure 4A)**. All accessions within the main Group 2 were assigned to the sub-populations 2, 3 and 5, while sub-populations 1, 4, 6-8 were specific to Group 1.

**Figure 4:**
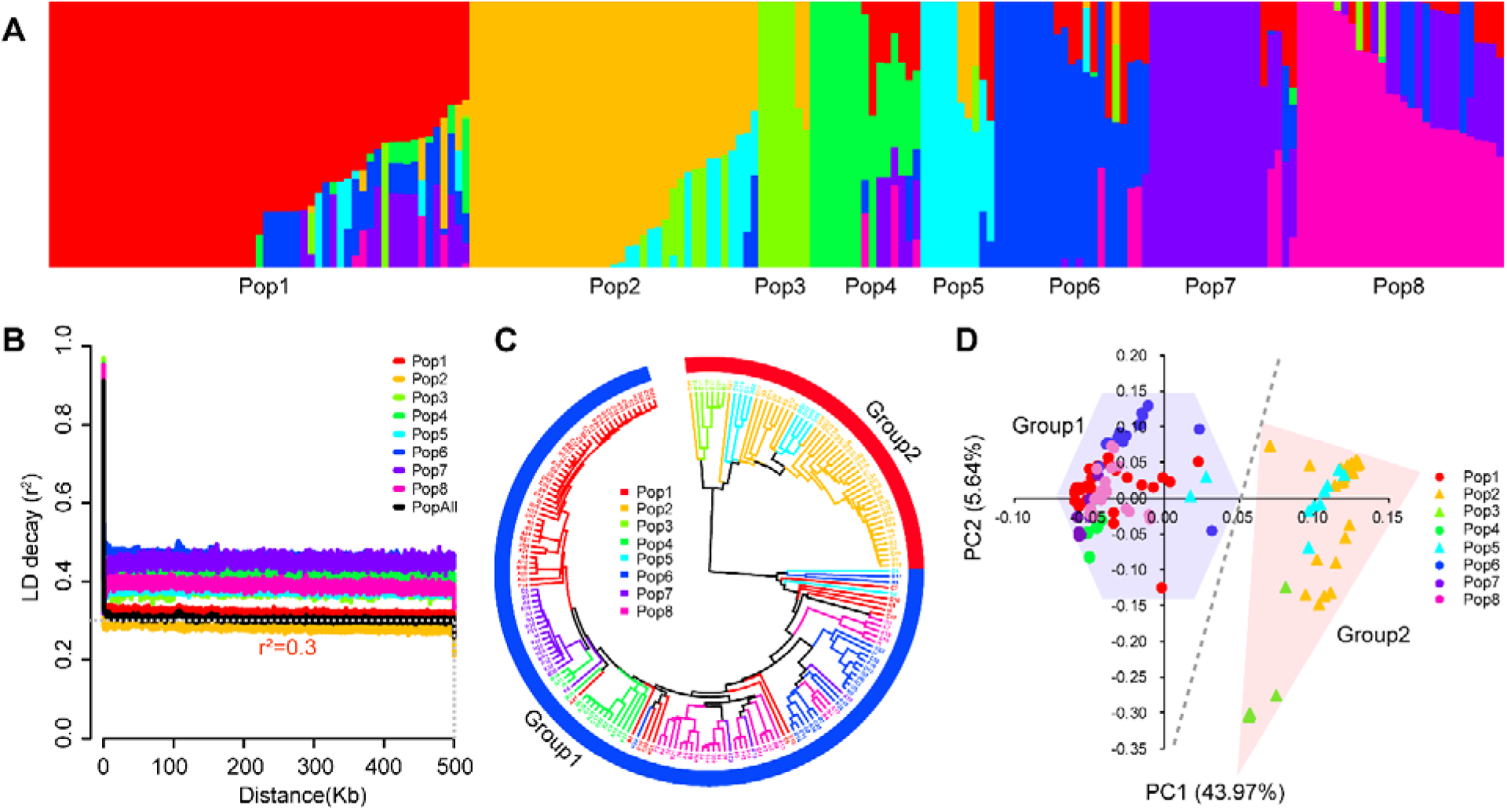
Population structure of 197 groundnut varieties. **A** Based on the computation results of fastStructure, 197 varieties have been classified into 8 subgroups. **B** Genome-wide averaged distance of LD decayed for different subgroups. **C** Phylogenetic tree of all varieties. Correspondences between branches and main subgroups from fastStructure are indicated by colors. Eight subgroups were merged into two groups. **D** PCA plot of the first two components for all varieties. The dot colors correspond to the 8 subgroups. Different shape coverage areas represent different groups.

The PCA plot revealed that the first two principal components (PCs), accounted for 49.61% of the total genetic variation **(Figure 4D)**. The broad distribution of accessions along the PC axes indicated a high genetic diversity within the panel. The phylogenetic tree further supported the population structure analysis, consistently resolving two main groups with eight subpopulations. Notably, the two groups exhibited distinct phenotypic profiles **(Figure 5).** Group 1 was distinguished by significantly higher values for yield related traits, like 100-kernel weight, kernel width, kernel length, 20-pod length, and overall yield. While Group 2 had significantly higher values of plant architecture related traits plant height, stem girth and leaflet length and width **(Figure 5A, D, E and F)**. Traits like plant width, shelling percentage and number of branches did not differ significantly between the two groups **(Figure 5B, C and I)**. The extent of linkage disequilibrium (LD), estimated by r^2^, varied among subpopulations. Genome-wide analysis revealed that the average distance at which LD decayed to a threshold of r^2^ = 0.3 was approximately 250kb across subpopulations. Subpopulation 6 exhibited the slowest LD decay, followed by sub-population 7, whereas subpopulation 2 showed the fastest LD decay **(Figure 4B)**.

**Figure 5:**
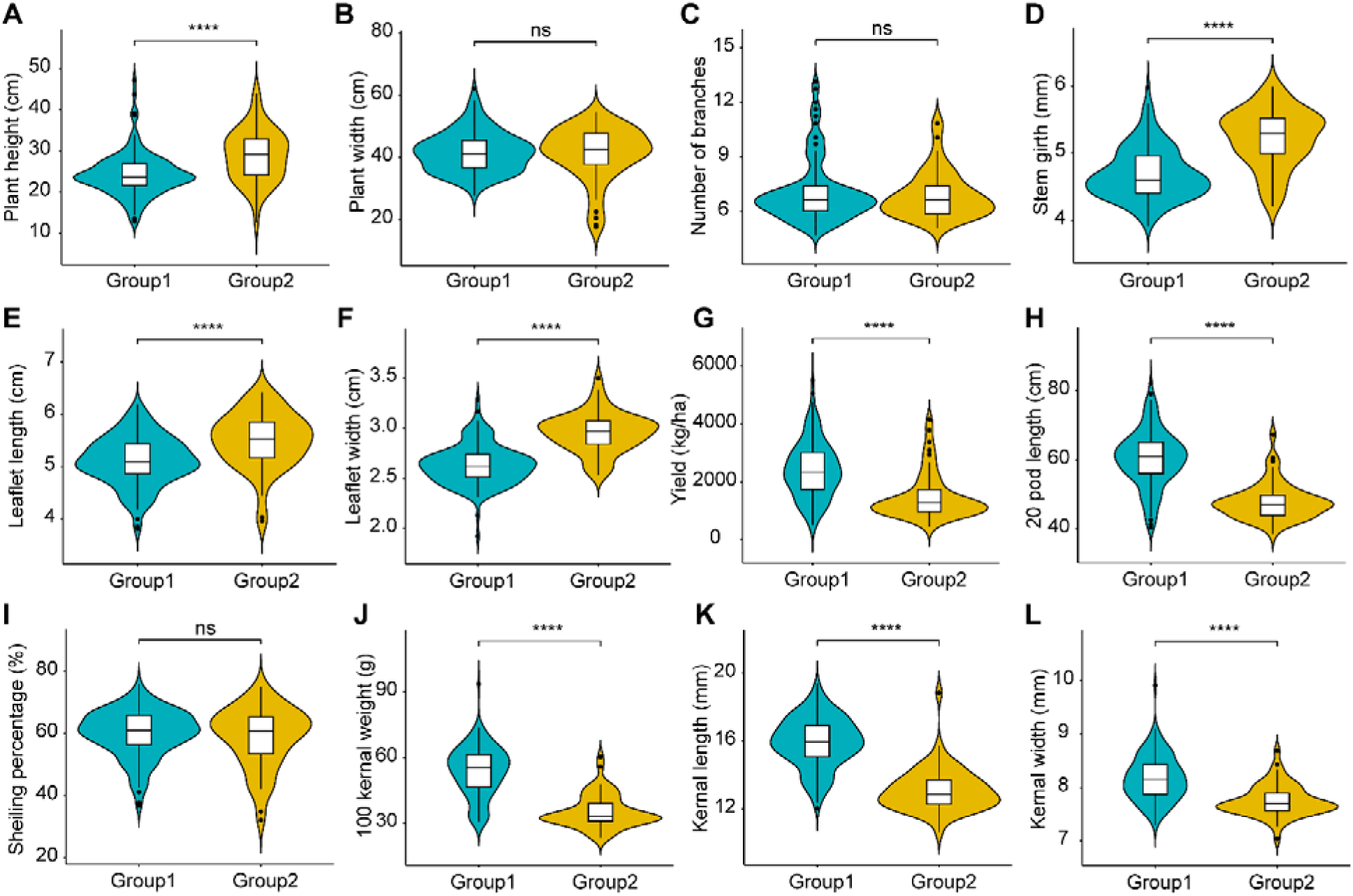
Comparison of phenotypic differences between groups. **A-L** Comparison of differences between groups in different traits based on t-test. **** indicates that *P* < 0.0001, ns represents that *P* > 0.05.

### Signatures of local adaptability and selective sweeps

The cross-population composite likelihood ratio test (XP-CLR) identified loci under divergent selection by comparing two distinct groups in the diversity panel. A total of 85 selective sweeps were identified, with XP-CLR scores ranging from 53.5 to 435.7. To minimize false positives, only top 1% of the most significant selective sweeps are discussed here **(Supplementary Table S5 and Figure 6)**. Several strong selection signatures were detected between the two groundnut groups on chr1, chr2, chr5, chr12, chr17, chr18 and chr19. On chr1, a genomic region spanning 5.5 to 6.01 Mb (containing 70 genes) was under strong selection pressure. The peak SNP within this region was located in a gene encoding a trihelix transcription factor (GT-3b-like) and an autophagy related protein 9-like gene. GO analysis

**Figure 6.**
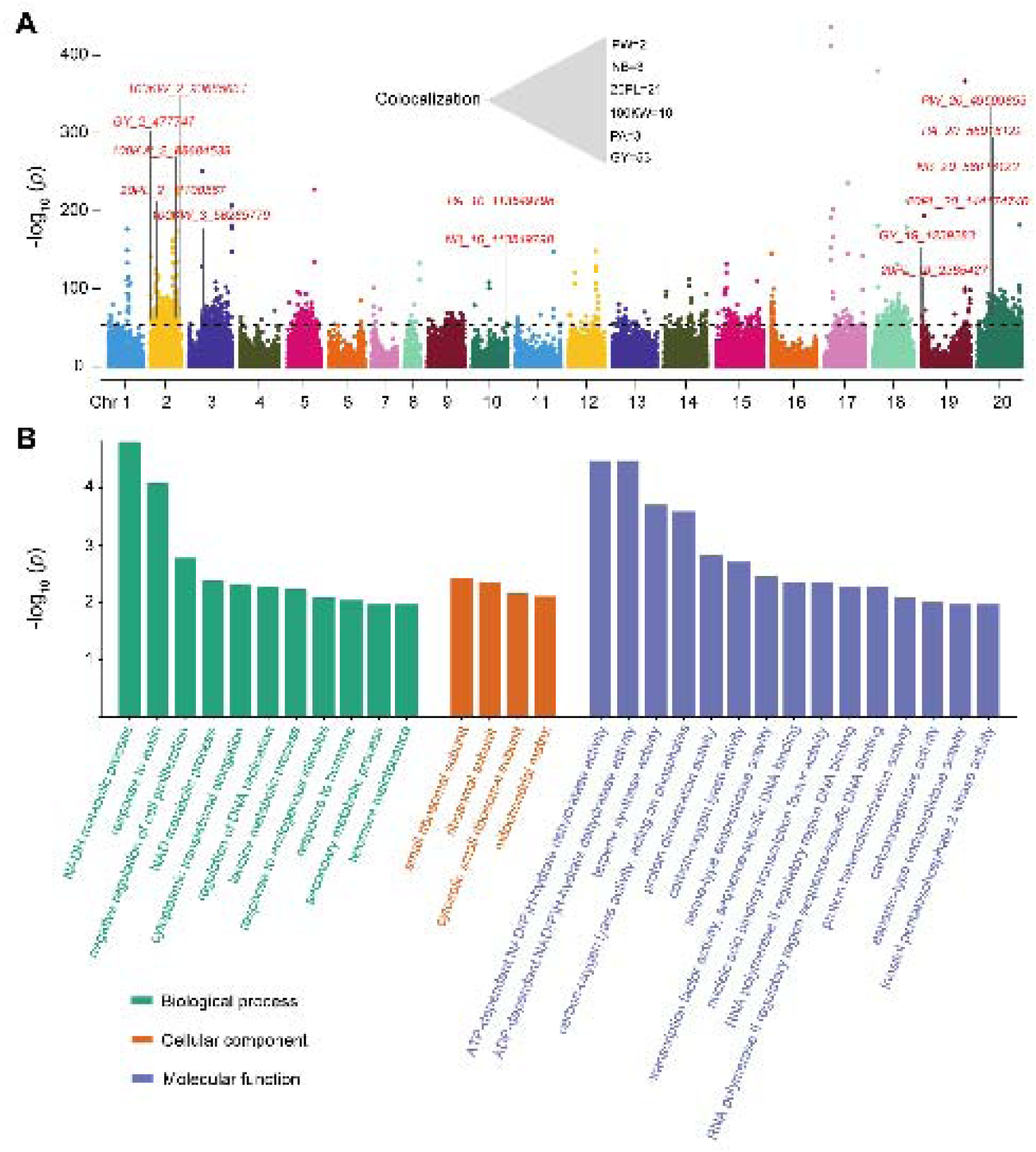
Profiling of the selective sweeps between two groups. **A** Genome-wide selective signals (XP-CLR score) between group1 and group2. The black dashed lines represent the cutoffs that define statistical significance (53.52). The GWAS loci mapped in this study are marked above the selective signal peaks. **B** GO enrichment analysis of genes in selective sweeps regions. The most significant top 30 GO terms are displayed.

Likewise, another region on chr1 (62.1 to 62.4), consisting of 45 genes has an XP-CLR score exceeding 100. The peak SNPs within this region were located in genes encoding several F-box family proteins and senescence associated proteins, suggesting a role in growth regulation and programmed cell death. The highest XP-CLR score was obsereved for a 0.3 Mb region at the distal end of chr17 (20.5 to 20.8 Mb) containing 10 protein encoding genes. Among these, four genes encoded cytochrome P450 superfamily proteins, and two genes encoded MADS-box and MYB transcription factors, both of which are known to regulate the expression of key genes involved in…………………………..

### GWAS for yield and plant architecture related traits

GWAS using whole-genome resequencing data identified 60 loci associated with 12 aforementioned traits. The maximum number of loci (n=20) were identified for number of branches and 20-pod length **(Supplementary Table S6 and S7; Figure 7)**. Only two loci were identified for yield and kernel length. while no loci reached the designated significance threshold for leaflet width and leaflet length. For plant height, five loci were most significant, located on chr11 (56.1 and 138.9 Mb), chr14 (9.7 Mb), and chr16 (5.5 and 9.3 Mb). However, LD analysis between the two SNPs on chr16 (5.5 and 9.3 Mb) showed that both loci were linked. There were 4 loci associated to plant width, with the most significant locus located on chr20 at 25.1 Mb **(Supplementary Table S6)**. Similarly, twelve marker traits associations were identified for number of branches (NOB) with the most significant SNP on chr10 (positioned at 113129995) with the *p* value of 9.32E-10. Two loci were associated with stem girth, the most significant of which was located on chr4 (positioned at 21698759, *p* ≤ 9.91E-07). Another peak associated with groundnut architectural traits was identified on chr9 with a tag SNP at position 4949939 bp **(Figure 8A)**. For yield two significant loci were identified on chr2 (positioned at 4969793 bp) with the *p* value of 2.82E-07. For 20-pod length a total of twelve marker traits associations were identified, with the most significant SNP located on chr12 (positioned at 8220852 bp, *p* ≤ 8.79E-12) **(Figure 8B)**. Three marker traits associations were detected for shelling percentage on chr19 (positioned at 26282634 bp, *p* ≤ 2.46E-07). Similarly, three marker traits associations were identified for 100-kernel weight on chr3 (positioned at 44863205 bp, *p*-value (8.37E-07). A total of four marker traits associations, two each on chr19 (positioned at 155786307 bp with a *p* value of 4.63E-06) and kernel width on chr7 (positioned at 36211773 bp) with a *p* value of 5.46E-08.

**Figure 7.**
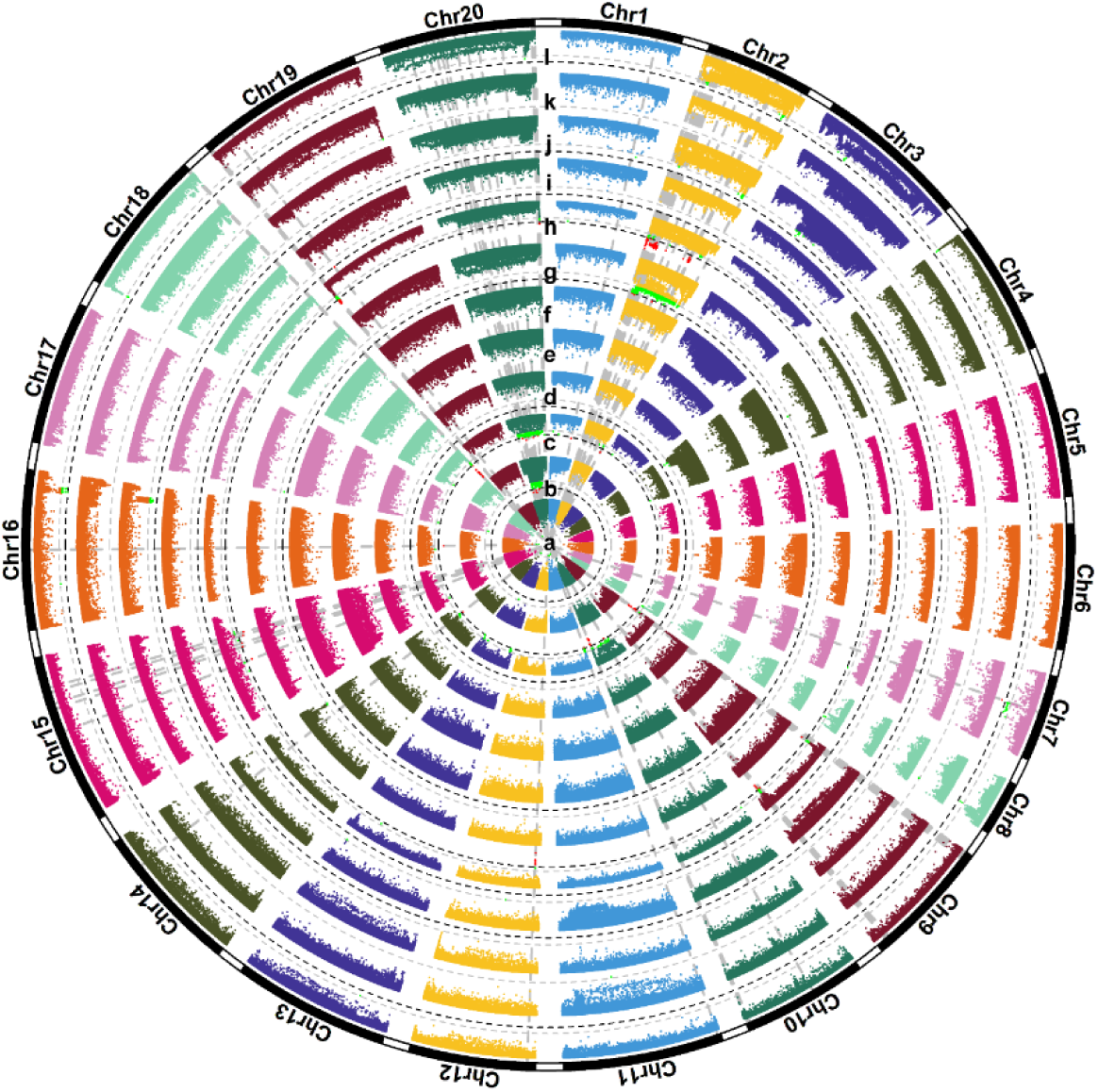
Manhattan plots of GWAS results for 12 traits. **(a-l)** represent plant height, plant width, number of branches, stem girth, leaflet length, leaflet width, yield, 20 pod length, shelling percentage, 100 kernel weight, kernel length and kernel width, respectively. The gray and black dashed lines represent the recommended threshold (6.10E-6) and significance threshold (3.05E-7), respectively.

**Figure 8.**
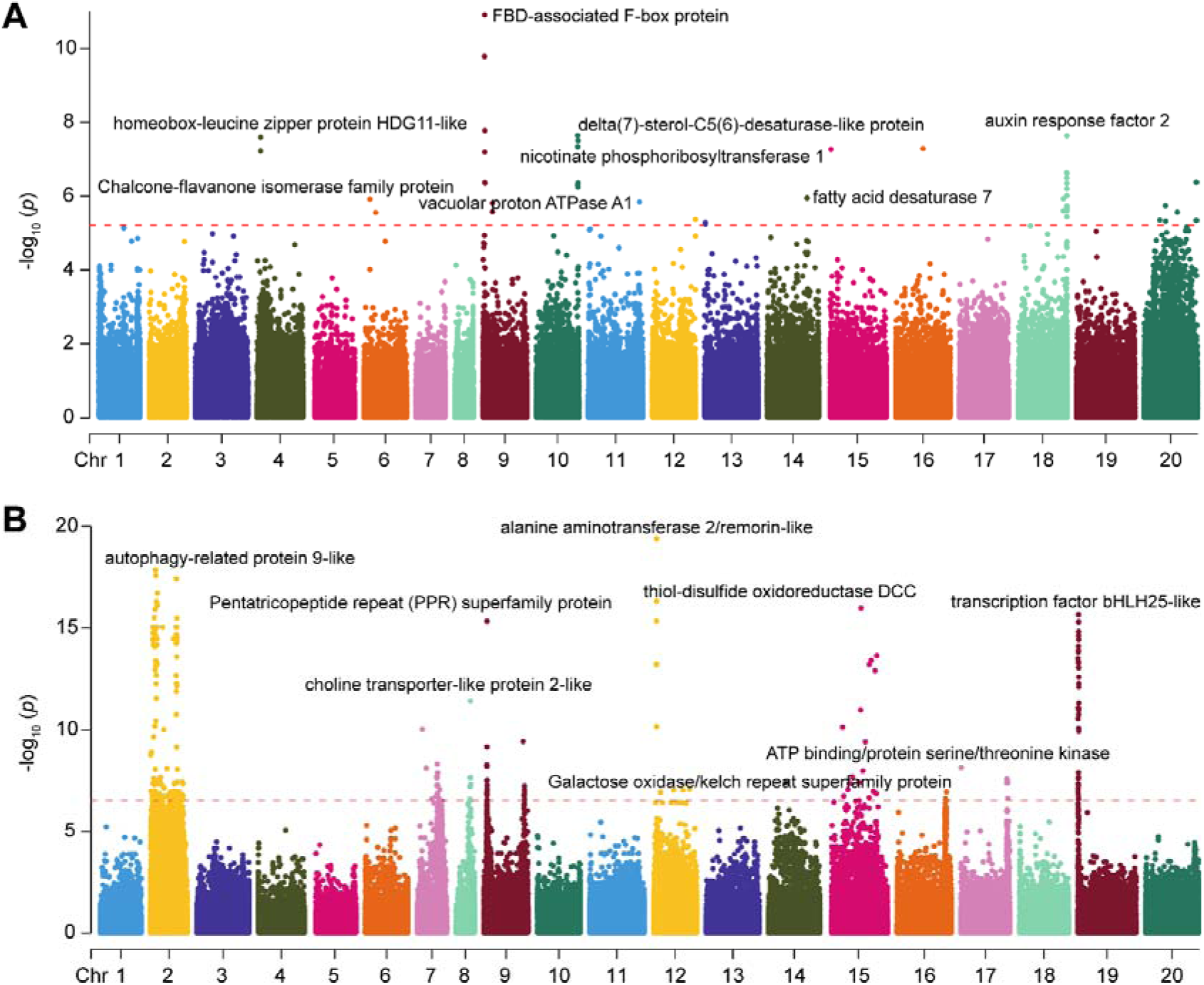
Manhattan plots of multiple trait GWAS. **A** Manhattan plot of GWAS result for plant architecture related traits. The red dashed lines represent the recommended threshold (6.10E-6). **B** Manhattan plot of GWAS result for grain yield related traits. The red dashed lines represent the significance threshold (3.05E-7). The candidate gene closest to peak SNPs are marked.

### Co-segregation of traits associated loci under selective sweeps

To confirm the selection signals identified by FST analysis, an allele-frequency based approach was applied using the cross-population composite likelihood ratio test (XP-CLR) as it is not affected by ascertainment bias and offers the advantage of signal enhancement. A total of 92 co-localizations were identified between GWAS signals and genome wide selective sweeps **(Table S8)**. For pod width, only two co-localizations were found, both on chr20, corresponding to positions PW_20_Fortnite 509855 and BMW_20_79748665 with XP CLR scores of 68.76 and 70.50 respectively. Three co-localizations were found for NOB one on chr10 (positioning 113549798) and two on chr20 positioning (42653358 and 58018122) respectively, with XP CLR scores of 56.04, 54.39 and 66.61, respectively. Similarly, a total of 21 co-localizations were found in selective sweeps and GWAS results for 20-pod length. The highest XP-CLR Score (106.19) was observed on chr2 at position 16103863 bp, while the remaining 20 co-localizations were distributed across chr2 and chr20 with XP-CLR scores ranging from 106.19 to 53.60. For HKW, 10 co-localizations were found with the highest XP-CLR score of 79.81 on chr2 at position 88684539 bp. The remaining 9 co-localizations were located on chr2, chr3 and chr20 with XP-CLR scores ranging from 79.81 to 55.40.

Further plant architecture traits we identified a total of 3 co-localizations with the highest XP-CLR score of 66.61 on chr20 at ∼58018122 bp. The other two co-localizations were found on chr10 and chr20 with XP-CLR scores of 56.04 and 54.39, respectively. For yield related traits, 53 co-localizations were identified, with the highest XP-CLR score of 114.65 located on chr2 at ∼83140648 bp. The remaining co-localizations were distributed across chr2 and chr19 with XP-CLR scores ranging from114.65 to 53.604.

### Cross validation of the chr9 locus associated with plant architecture traits in the groundnut355 panel

A regional association plot on chr9 was generated for plant architecture traits using GWAS **(Figure 9A)**. The most significant peak in the present study was identified on chr9, representing an association with a cluster of FBD associated F-box protein genes. This genomic region spans from 4.450 Mb to 5.447 Mb covering 996.76 kb, with a significance threshold of 3.05E-7. The strongest association signal was observed at SNP position 4949939 bp (*p* = 3.6E-10), with the putative candidate gene, *Arahy.37HYKA*. This LD block was associated with several plant architectural traits including plant height, number of branches, stem girth, leaflet length and width **(Figure 9B-G)**. To validate these findings, we mined the groundnut355 resequencing data to identify corresponding SNPs associated with plant architectural traits. LD analysis of SNPs in the groundnut355 panel revealed that a haplotype block, hap09-5368928_5468037, was in perfect LD (r^2^=1) with the tag SNPs in our diversity panel on chr9 at this locus. Within this haplotype block, three genes were annotated; *Arahy.EPZV43* (ATP-binding ABC transporter), *Arahy.550YKG* (CDPK-related kinase), and *Arahy.9VTD7S* (ABC transporter family). The haplotype frequencies were examined across subspecies **(Figure 9H)**. Hap-IV was only present in *A. hypogeae* irregular-hypogeae type, while Hap-I was in subspecies fastigiate var vulgaris.

**Figure 9.**
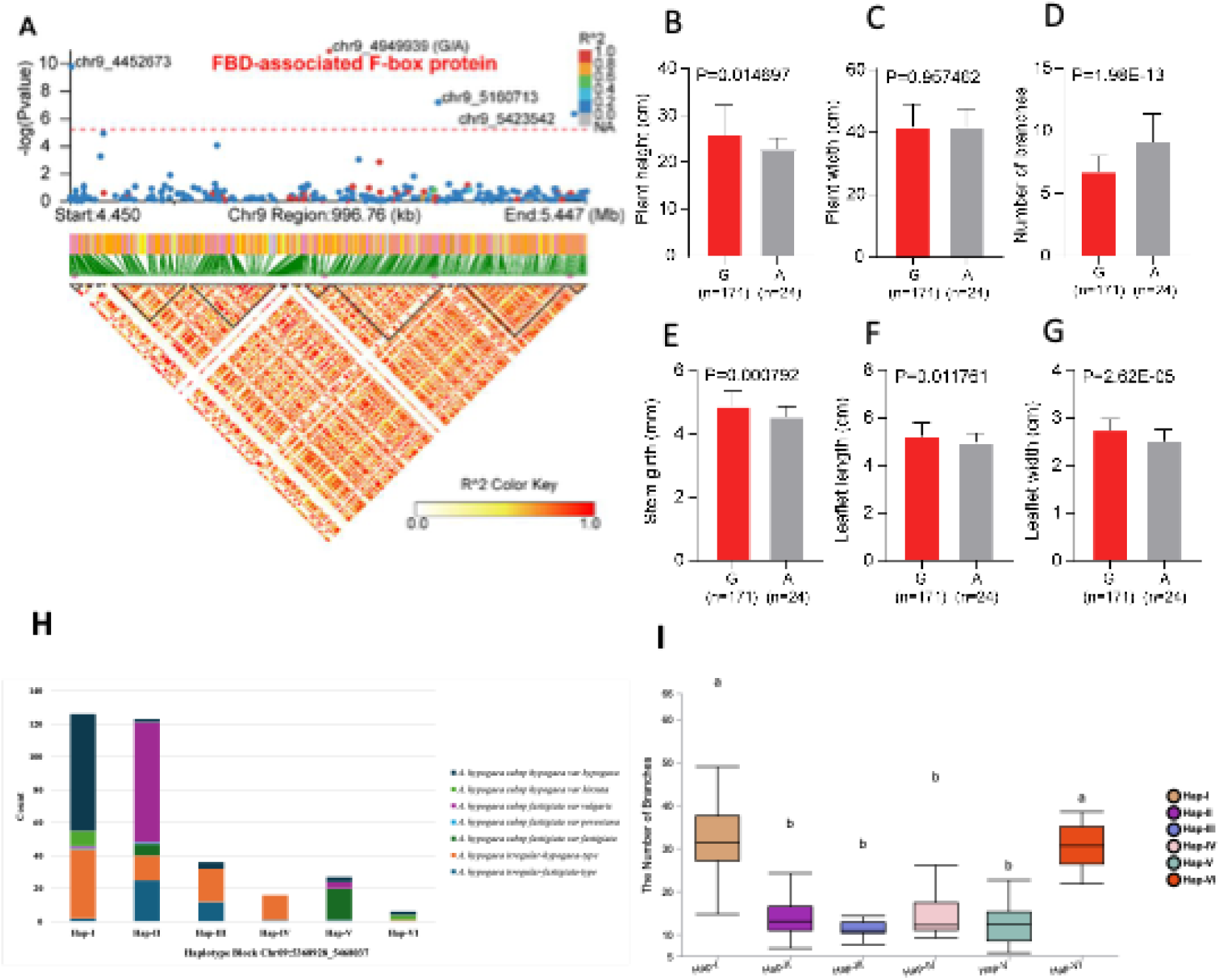
Regional association plot and effects of alleles and haplotypes in groundnut diversity panel. **A** Regional association plots on chr9 for plant architecture related traits. The red dashed lines represent the recommended threshold (6.10E-6). **B** Regional association plots on Chromosome 12 for grain yield related traits. The red dashed lines represent the significance threshold (3.05E-7). **C-G** Comparison of differences in different haplotypes for different traits. **H** Haplotype distribution of Haplotype block chr9:5368928-5468037 on chr9 in groundnut355 diversity panel in different ecological types of *A. hypogaea*. **I** Phenotypic effects of six haplotypes on the number of branches, the boxplots sharing the same letters are not significantly different at *P*=0.01.

The haplotypes showed significant association with number of branches in the groundnut355 panel, with Hap-I associated with a significantly higher number of branches compared to other haplotypes, while Hap-III had the least number of branches **(Figure 9I)**. The expression profiles of key genes within this haplotype were extracted from Sinha et al (2020), and only two genes, *Arahy.EPZV43* and *Arahy.550YKG*, showed detectable expression profiles. *Arahy.550YKG* was expressed only in seeds, nodules, and stems, while *Arahy.EPZV43* showed expression across all tissues, with the highest expression in seeds and nodules.

### Cross validation of chr12 locus associated with multiple yield related traits and validation in groundnut355 panel

A regional association plot on chr12 was generated for yield related traits using GWAS **(Figure 10A)**. The peak SNPs were distributed between 7.72 Mb to 8.714 Mb, spanning 992.03 kb, with a significance threshold of 6.10E-6. The strongest association signal was observed at SNP 8221397 within *Arahy.E9MTVL*, a gene encoding an Alanine aminotransferase remorin/like protein, with a *p*-value of 8.79E-12. This genomic region and its peak SNPs were strongly associated with multiple yield related traits including Yld, 20-pod length, shelling percentage and HKW **(Figure 10B-G)**. The corresponding genomic region was extracted from the groundnut355 panel and haplotype analysis confirmed that a haplotype block in the same region was associated with yield related traits. Two haplotypes Hap-II and Hap-III, restricted to *A. hypogaea* var. vulgaris and var. fastigitata **(Figure 10H)** and were associated with lower yield related trait values. However, Hap-I, which was restricted to the *A. hypogaea* irregular-hypogaea-type subspecies, was associated with higher seed length, HPW and HKW in the groundnut355 panel. Given that this region has multiple genes in strong LD, further investigations are required to identify the causal genes within this region and elucidate their functional roles in regulating peanut yield.

**Figure 10.**
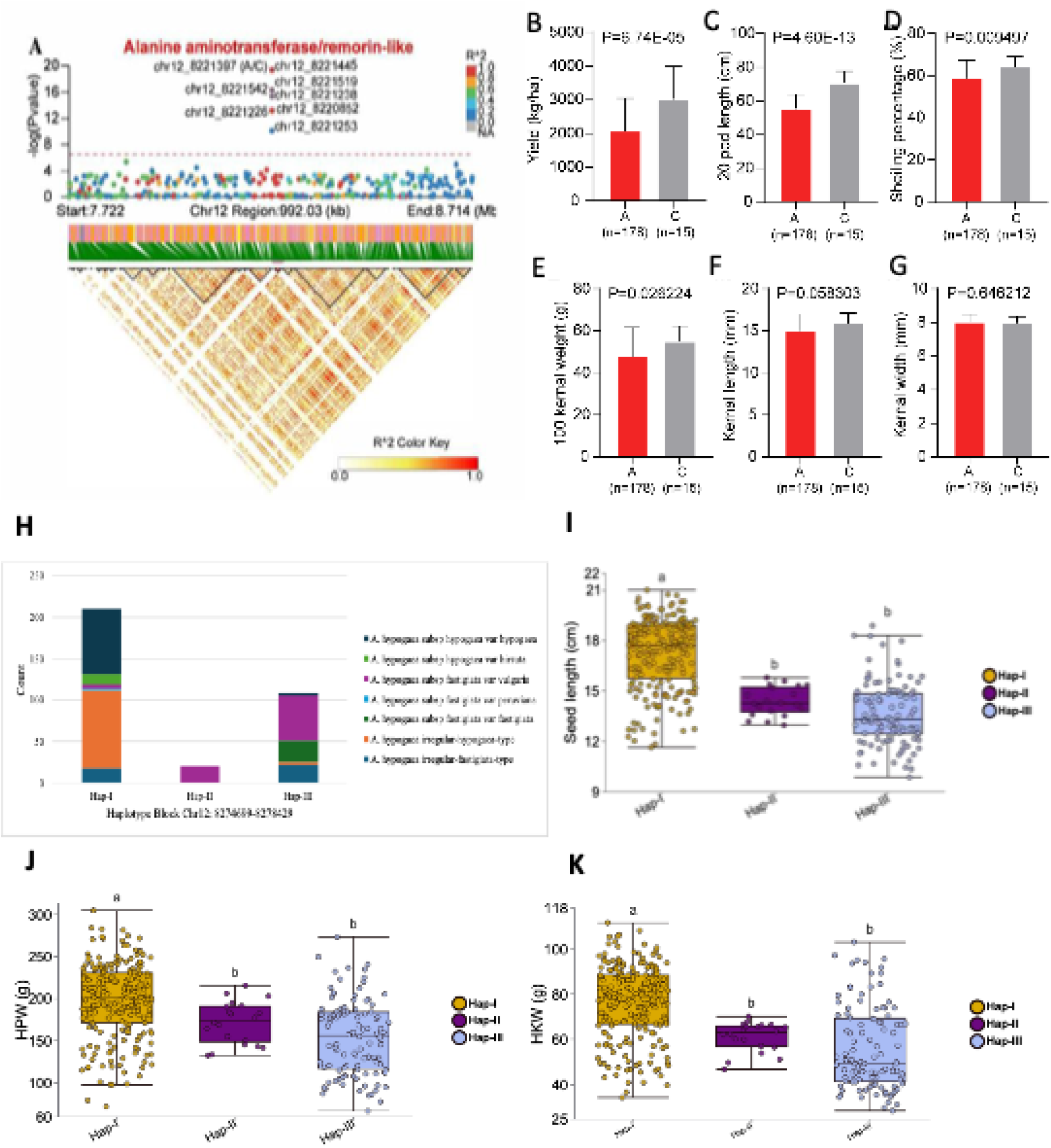
Regional association plot and effects of alleles and haplotypes in groundnut diversity panel. **A** Regional association plots on chr12 for yield-related traits. The red dashed lines represent the recommended threshold (6.10E-6). **B-G** Comparison of differences in different alleles of the tag SNP associated with multiple yield-related traits. **H** Haplotype distribution of Haplotype block chr12:8274699-82778429 on chr12 in groundnut355 diversity panel in different ecological types of *A. hypogaea*. **I-K** Phenotypic effects of three haplotypes on seed length, hundred pod weight, and hundred kernel weight; the boxplots sharing the same letters are not significantly different at *P*=0.01

## Discussion

### Phenotypic variation in groundnut diversity panel

In the present study a groundnut diversity panel was assembled and characterized for yield and plant architecture traits. The phenotypic variation in the diversity panel was more pronounced in yield, 100-kernel weight and 20-pod length. This can be mainly attributed to the diverse geographic origin of the groundnut population. A global collection of 390 groundnut accessions collected from China, USA and India was characterized for 28 agronomic traits and substantial amount of phenotypic diversity was observed (Lu et al. 2024). The geographic background is a key factor in delivering morphological variation for economically important vital traits such as pod yield, grain yield, number of pods per plant, and oleic acid percentage. Our diversity panel showed near normal like distribution for plant height, plant width and leaflet length which demonstrates potential of utilization of this material while breeding for plant architecture and canopy structure. Broadly all the quantitative traits showed divergence in the population except for kernel width, stem girth and leaflet length. Banla et al. (2020) justified the lesser extent of variation in these quantitative traits because of a directional selection which reduced cover-wide phenotypic variance, and the results were also supported in other studies (Upadhyaya et al. 2003). Previously studies have reported significantly lower yield for groundnut cultivars in Pakistan where in some cases the yield obtained is nearly half as low as that of major groundnut growing countries (Jahanzaib et al. 2019). Nearly a fifth of the diversity panel reported mean yield of more than 3000 kg ha^-1^ which makes these accessions as suitable candidates for selection under target environmental conditions as high yielding cultivars. The Pearson’s correlation showed strong positive associations between various yield related traits, whereas Yld negatively correlated with stem girth, leaflet width, and plant height. The characteristics associated with grain size and shape play a direct role in determining yield, with these traits collectively regulated by both maternal and nuclear genes. This implies the potential for their concurrent selection (Venuprasad et al. 2011). The vegetative traits such as number of branches positively contributed towards the grain yield in groundnut germplasm (Amoah et al. 2023).

### Genomic diversity, population structure and LD decay

The genome variation map developed for our diversity panel showed high SNP density across genome with variable number of SNPs on different chromosomes. For example, the highest density of 350 SNPs/Mb (3.5 SNPs/kb) was observed on chr16 whereas SNPs were not dense for chr1 and chr8. Also, there were 1.4-fold greater number of SNPs on B subgenome (∼0.3 million SNPs) compared to A subgenome (0.41 million SNPs). Similar results were reported earlier and 1.36-fold number of SNPs were observed in a global groundnut collection (Lu et al. 2024). The higher SNPs density rate at certain chromosomes like chr16 suggests potential hotspots of recombination or historical selection pressure (Renaut et al. 2014). Lai et al. (2015)Lai et al, (2014) attributed variable number of SNPs on different chromosome in wheat because of gene flow and evolution because of domestication and rigorous breeding.

The lower mean heterozygosity rate was anticipated due to self-pollinated nature of groundnut. However, parallel lower SNPs heterozygosity rate of 0.04 was recently reported in Kirstings groundnut (Akohoue et al. 2020) suggesting narrow genetic base of the studied population. These findings agreed with the previous report on groundnut SNPs variation (Bertioli et al. 2019; Sun et al. 2022).

FastStructure elucidated a total of two major groups which can be further divided into eight sub-populations. On the contrary recent molecular studies on groundnut populations have reported relatively small genetic structure with two (Bhawar et al. 2020), three (Abady et al. 2021) four or five sub-populations (Otyama et al. 2020). These studies have examined comparatively less diverse regional or interrelated populations suggesting that genetic structures get complex as the size of populations increases. This was further confirmed by (Kafoutchoni et al. 2021) as they observed eight sub-groups in the population comprising of 217 Kersting’s groundnut genotypes. The subsequent phylogenetic analysis of the genomic data revealed that the grouping was in harmony with phenotypic characteristics of the individuals. For example, the first major group comprising of subgroup-I, II, V and VI was characterized with higher oil content, kernel width and shelling percentage whereas the second group housed subgroup-III, IV, VII and VIII which fared low for these traits. The population structure in plants and animals shape the phenotypic variation and likely the natural selection is a major force driving population differentiation (Pokrovac and Pezer 2022).

LD decay was estimated across population and within subpopulations. The higher LD decay rate in subpopulation 6 showed that individuals tend to be more homozygous for various alleles, and individuals in this subpopulation included approved cultivar (BARD-19) and advanced lines from Pakistan. Conversely the lower LD decay rate in subpopulation 2 consisting of diverse lines from different countries indicated the somewhat frequent shuffling of the alleles during recombination. This could potentially be explained by the possibility that the group represents an admixed and outbred population. LD decay is function of size of population and genomic regions, hence variable LD decays rates have been reported for different crops by different studies (Li et al. 2022) (Rafalski and Ananiev 2009). For example, maize shows rapid LD decay after 5–10 kb, but in certain regions of the genome, LD can last hundreds of kb (Andrade et al. 2019). This was confirmed in groundnut as they discovered rapid decline in LD with distance, in the ICRISAT groundnut mini-core collection, with LD dropping below 0.2 at 41.6 kb (Shaibu et al. 2020). The recent whole-genome resequencing of global groundnut population showed LD decay 92.3 kb (r^2^ ∼0.16) (Lu et al. 2024), and LD decay distance was 240 kb in our diversity panel for r^2^=0.3 which is comparable.

### Genes underpinning local adaptability

Modern crop cultivars are consequence of artificial selection for yield and quality traits which resulted in the dramatic improvement of many agronomic traits. However, it has resulted in narrow genetic diversity generally in food crops. Therefore, it is important to assess the selection pressure on specific loci to identify the opportunities for future crop breeding. We identified 4525 SNPs within selective sweeps spanning the whole genome. The distribution of sweeps across A and B genome was almost symmetrical as the number of sweeps for both types was 2348 and 2177, respectively. The highest number of selective sweeps between two groups were observed on four chromosomes i.e. chr20, chr18, chr16 and chr19 exhibiting strong GWAS signals for yield and plant architecture traits. Notably, on chr2 and chr19, SNPs linked with 20-pod length, 100 kernel weight, and grain yield co-localized, indicating an important instance of selective sweep primarily driven by selection pressure aimed at developing high-yielding varieties over a century-long period (Gangurde et al. 2020).

The plant architecture traits such as plant width and number of branches determine the canopy size and growth habit of the groundnut. On chr10, a locus represented by tag SNP 10_113549798 was associated with number of branches and plant architecture traits in single-trait and multi-trait GWAS, respectively. Also, four SNPs strongly associated with both yield (pod width and 20 pod length) and plant architecture traits on chr20, exhibited selective sweep signatures, surpassing the threshold XP-CLR score, suggesting their co-localization as a result of selection pressure. This phenomenon can be also explained by the fact that effective population size of the groundnut has seen a dramatic expansion during ca 300-500 years (Liu et al. 2022). Selective sweeps linked to QTL and genes governing pod and seed size, plant architecture, and/ or disease resistance align with the primary objectives of enhancing groundnut breeding. A GO enrichment analysis of the candidate genes in the selective sweep region exhibited the top thirty GO terms having role in the biological processes, molecular function or acted as a cellular component. Notably candidate genes in these regions regulated functions such as NADH metabolic processes, response to auxin, regulation of DNA replication, Leucine metabolic process, synthesis of small ribosomal subunits and range of molecular activities. Most of these candidate genes have been recently reported by researchers tracing groundnut domestication, evolutionary biology and selection pressures post breeding era (Liu et al. 2022; Wang et al. 2019; Zhang et al. 2017). These genes either directly or indirectly play their role in development of plant seed and architecture as well as response to environmental stimuli for better adaptation (Mallikarjuna and Varshney 2014).

### Discovery of trait associated loci and validation

A large number of SNPs (9844) were associated with agronomic traits likely due to the use of more number of quantitative phenotypes. These SNPs were narrowed down based on the genome positions and were restricted to 60 high-confidence loci. Wang et al. (2019) carried out GWAS to determine the genomic basis of phenotypic variation in seven yield related traits and identified a total of 26 significant peak SNPs for primary branches per plant on five chromosomes. We reported 12 loci strongly associated with number of branches. Zhang et al. (2023) reported pod length related SNPs on chr1, chr2, chr3, chr4, chr7, chr10, chr12, chr13, chr14 and chr18 which was confirmed by our findings for 20-pod length. The QTL/GWAS database at www.peanutbase.org was mined for the co-localized SNPs, and we found six SNPs within the genomic regions reported earlier for pod-related morphological traits. These SNPs include loci on chr2 at 6.6 Mb and 54.8 Mb, chr9 at 8.3 Mb, chr15 at 99.2 Mb and chr17 at 125.6 and 128.3 Mb (Gangurde et al. 2020). For pod width, the significant association with eight SNPs distributed on chr3, chr4, chr7, chr13, chr16, chr18 and chr20 in this study were in congruence with findings of the previous reports (Gangurde et al. 2020; Zhang et al. 2023; Zhou et al. 2021).

Yield is a complex trait mainly augmented by the additive effects of both architectural and seed related traits. In the current study phenotypic variability for grain yield depicted strong association with two SNPs localized on chr2 at 9.7 Mb and 43.5 Mb. In the literature SNPs have been localized on chr5, chr6, chr8, chr14, chr15, chr16 and chr18 showing strong association with yield related traits such as seed and pod weight. Contrastingly, these chromosomes did not harbor any of the yield linked SNPs in the current investigation, moreover, the pod width and length related SNPs did co-localized with those reported by previous studies (Gangurde et al. 2020; Luo et al. 2017).

Plant height is an important architectural trait controlling yield, economic value and harvest index due to its strong effect on light interception and photosynthesis (Sarlikioti et al. 2011). In the current study, five SNPs located on chr11, chr14, chr16 and chr19 showed significantly strong association with plant height. A recent meta-study on elucidating genetic basis of plant height in groundnut detected strong signals on chr11, chr14, chr16 and chr19 (Sahu et al. 2025). Huang et al. (2016) constructed a high-density genetic map of groundnuts using SSR markers and transposon-based techniques in a recombinant inbred line (RIL) population. Their study revealed the presence of 18 QTL, with 7 located on chr4, chr5, chr13 and chr16. These findings diverge from the outcomes of our present investigation, likely stemming from disparities in the marker systems employed between the two analyses.

### Candidate genes for plant architecture and yield-related traits

Within the 1Mb of suggestive and significantly associated SNPs for both plant architectural traits and yield related traits 16 candidate genes were identified. SNPs showing strong associations with plant architecture related traits were co-localized with eight annotated genes such as FBD-associated F-Box protein, homeobox-leucine zipper proteinHDG11-like, delta(7)-sterol-C5(6)-desaturase-like protein, auxin-response factor 2, nicotinate phosphoribosyltransferase 1, chalcone-flavonone isomerase family protein, fatty acid desaturase 7 and vacuolar proton ATPase 7. In a genomic region spanning 996.76 kb, located between 4.4 to 5.44 Mb on chr9, four significant SNP signals associated with primary branches per plant and 20-pod length were identified. Notably, one of these SNPs, positioned at 4949939 bp, was found to be adjacent to an FBD-associated F-box protein. A recent study has confirmed the role of F-box protein in groundnut lateral branch development (Li et al. 2024). F-Box protein family is the one of the largest studied in plants which are encoded by genes responsible for plant growth and development, photomorphogenesis, plant leaf morphology and photopigments (Feng et al. 2023; Mo et al. 2021). Li et al. (2022) utilized GWAS and bulk segregant analysis to investigate the genetic loci controlling traits associated with growth habit in groundnut. Their analysis unveiled significantly strong associations between SNPs and growth habit traits, leading to the identification of 597 candidate genes. Notably, among these candidates, 29 genes were found to encode the F-box protein, emphasizing its involvement in cellular protein degradation. Likewise, Zhang et al. (2017) while investigating the genomic background of pod related traits, identified one candidate gene of F-box protein in addition to bHLH transcription factor, major intrinsic protein (MIP) and proline rich protein (PRH) genes.

A regional association plot depicting a 992.03 kb genomic region revealed a highly significant strong association at chr12 for 20-pod length which is a major yield contributing trait. This loci colocalized with Alanine aminotransferase 2 (AlaAT2)/ remorin-like gene. Other important genes in the vicinity of the region included autophagy-related protein 9-like, pentatricopeptide repeat (PPR) super family protein, thiol-disulphide oxidoreductase (DCC), transcription factor bHLH25-like, choline transporter-like protein 2-like, ATP binding/proteinserine/threonine kinase and Galactose/oxidase/ kelch repeat super family protein. Alanine aminotransferase 2 (AlaAT2) is an important plant enzyme playing its role in nitrogen metabolism. Research has confirmed the significance of AlaAT enzymes in amino acid synthesis pathways, thereby enhancing nitrogen utilization efficiency (NUE) in plants (Jiang et al. 2013; McAllister and Good 2015). These enzymes, including AlaAT2 homologs found in plant peroxisomal glyoxylate aminotransferases, are integral components of the photorespiratory pathway, underscoring their metabolic importance in plant physiology. Moreover, the enzymatic activity of alanine aminotransferase has been evaluated during the germination process in various crops revealing varying increases in activity across different germination stages. This underscores the enzyme’s significance in seedling development and resilience to climate extremes (Agrawal et al. 2024). Overall, AlaAT2 plays a critical role in nitrogen metabolism and amino acid synthesis pathways in plants, thereby influencing plant growth and development.

Conclusively, we dissected the important genomic regions underpinning economically important traits in groundnut with greater resolution and laid a strong foundation to delineate these genes for their function studies and breeding application. Two genomic regions were restricted to less than 1 Mb region with very few candidate genes underpinning groundnut developmental and yield potential.

## Conflict of interest

We declare no conflict of interest.

## Data availability

The whole genome resequencing data is deposited at NCBI project PRJNA1141749. The SNP variant file is available at https://zenodo.org/records/14388053.

## Funding

Authors thank for the financial support from Key R&D Programs of Hainan Province (ZDYF2024XDNY210), National Natural Science Foundation of China (32361143514), Project of Sanya Yazhou Bay Science and Technology City (SKIC-JYRC-2024-55), Nanfan special project, CAAS (YBXM2506), and the Agricultural Science and Technology Innovation Program (CAAS-CSIAF-202303). We also acknowledge Higher Education Commission (HEC) funding program NRPU-15269 for partial support.

## Authors contributions

MJ collected the data and analyzed the data. KH analyzed the data and wrote the first draft. SR analyzed the data. IK and HK collected the phenotypic data and contributed to draft writing. MJU analyzed the data and edited the manuscript. SG edited the manuscript. AR and HL designed the study, edited the manuscript and prepared the final draft.

## Supplementary information

Table S1. List of the accessions used in this study. The assignment of the accessions to any of the 2 main groups and 7 subpopulation and their phenotypic values across two years for 12 morphological and yield related traits are also provided.

Table S2. Analysis of variance (ANOVA) showing variation in phenotypic traits in groundnut diversity panel in the field trials conducted over two years

Table S3. Basic statistics of the raw sequencing data generated on each accession of diversity panel

Table S4. Number of SNP markers and their density on each chromosomes identified by whole-genome resequencing in the groundnut diversity panel

Table S5. Selective sweep regions between Group1 and Group2.

Table S6. GWAS results of 12 traits based on single traits analysis.

Table S7. GWAS results of multiple trait analysis.

Table S8. Colocalization of GWAS and selective sweep analysis.

## Notes

### Competing Interest Statement

The authors have declared no competing interest.

## References

Abady S, Shimelis H, Janila P, Yaduru S, Shayanowako AI, Deshmukh D, Chaudhari S, Manohar SS (2021) Assessment of the genetic diversity and population structure of groundnut germplasm collections using phenotypic traits and SNP markers: Implications for drought tolerance breeding. PloS One 16:e0259883

Afzal F, Li H, Gul A, Subhani A, Ali A, Mujeeb-Kazi A, Ogbonnaya F, Trethowan R, Xia X, He Z, Rasheed A (2019) Genome-Wide Analyses Reveal Footprints of Divergent Selection and Drought Adaptive Traits in Synthetic-Derived Wheats. G3: Genes|Genomes|Genetics 9:1957

Agrawal N, Chunletia RS, Badigannavar AM, Mondal S (2024) Role of alanine aminotransferase in crop resilience to climate change: a critical review. Physiology and Molecular Biology of Plants:1–19

Akohoue F, Achigan-Dako EG, Sneller C, Van Deynze A, Sibiya J (2020) Genetic diversity, SNP-trait associations and genomic selection accuracy in a west African collection of Kersting’s groundnut [Macrotyloma geocarpum (Harms) Maréchal & Baudet]. Plos one 15:e0234769

Amoah RA, Nelimor C, Gymafi BA, Boampong R, Osei CY, Yeboah A, Sackey V, Ansah EO, Awuah S, Mensah AO (2023) Analysis of agro-morphological variability and inter-trait relationships in Ghanaian groundnut (Arachis hypogaea L.) accessions. Plant Genetic Resources 21:471–479

Andrade ACB, Viana JMS, Pereira HD, Pinto VB, Fonseca e Silva F (2019) Linkage disequilibrium and haplotype block patterns in popcorn populations. PloS one 14:e0219417

Banla EM, Dzidzienyo DK, Diangar MM, Melomey LD, Offei SK, Tongoona P, Desmae H (2020) Molecular and phenotypic diversity of groundnut (Arachis hypogaea L.) cultivars in Togo. Physiology and Molecular Biology of Plants 26:1489–1504

Bertioli DJ, Cannon SB, Froenicke L, Huang G, Farmer AD, Cannon EK, Liu X, Gao D, Clevenger J, Dash S (2016) The genome sequences of Arachis duranensis and Arachis ipaensis, the diploid ancestors of cultivated peanut. Nature genetics 48:438–446

Bertioli DJ, Jenkins J, Clevenger J, Dudchenko O, Gao D, Seijo G, Leal-Bertioli SCM, Ren L, Farmer AD, Pandey MK, Samoluk SS, Abernathy B, Agarwal G, Ballen-Taborda C, Cameron C, Campbell J, Chavarro C, Chitikineni A, Chu Y, Dash S, El Baidouri M, Guo B, Huang W, Kim KD, Korani W, Lanciano S, Lui CG, Mirouze M, Moretzsohn MC, Pham M, Shin JH, Shirasawa K, Sinharoy S, Sreedasyam A, Weeks NT, Zhang X, Zheng Z, Sun Z, Froenicke L, Aiden EL, Michelmore R, Varshney RK, Holbrook CC, Cannon EKS, Scheffler BE, Grimwood J, Ozias-Akins P, Cannon SB, Jackson SA, Schmutz J (2019) The genome sequence of segmental allotetraploid peanut Arachis hypogaea. Nat Genet 51:877–884

Bhawar PC, Tiwari S, Tripathi M, Tomar R, Sikarwar R (2020) Screening of groundnut germplasm for foliar fungal diseases and population structure analysis using gene based SSR markers. Current Journal of Applied Science and Technology 39:75–84

Chen GB, Lee SH, Zhu ZX, Benyamin B, Robinson MR (2016a) EigenGWAS: finding loci under selection through genome-wide association studies of eigenvectors in structured populations. Heredity (Edinb) 117:51–61

Chen H, Patterson N, Reich D (2010) Population differentiation as a test for selective sweeps. Genome Res 20:393–402

Chen X, Li H, Pandey MK, Yang Q, Wang X, Garg V, Li H, Chi X, Doddamani D, Hong Y, Upadhyaya H, Guo H, Khan AW, Zhu F, Zhang X, Pan L, Pierce GJ, Zhou G, Krishnamohan KA, Chen M, Zhong N, Agarwal G, Li S, Chitikineni A, Zhang GQ, Sharma S, Chen N, Liu H, Janila P, Li S, Wang M, Wang T, Sun J, Li X, Li C, Wang M, Yu L, Wen S, Singh S, Yang Z, Zhao J, Zhang C, Yu Y, Bi J, Zhang X, Liu ZJ, Paterson AH, Wang S, Liang X, Varshney RK, Yu S (2016b) Draft genome of the peanut A-genome progenitor (Arachis duranensis) provides insights into geocarpy, oil biosynthesis, and allergens. P Natl Acad Sci USA 113:6785–6790

Chen X, Lu Q, Liu H, Zhang J, Hong Y, Lan H, Li H, Wang J, Liu H, Li S (2019) Sequencing of cultivated peanut, Arachis hypogaea, yields insights into genome evolution and oil improvement. Molecular plant 12:920–934

Danecek P, Auton A, Abecasis G, Albers CA, Banks E, DePristo MA, Handsaker RE, Lunter G, Marth GT, Sherry ST, McVean G, Durbin R, Genomes Project Analysis G (2011) The variant call format and VCFtools. Bioinformatics 27:2156–2158

Feng C-H, Niu M-X, Liu X, Bao Y, Liu S, Liu M, He F, Han S, Liu C, Wang H-L (2023) Genome-wide analysis of the FBA subfamily of the poplar F-box gene family and its role under drought stress. International Journal of Molecular Sciences 24:4823

Furlotte NA, Eskin E (2015) Efficient multiple-trait association and estimation of genetic correlation using the matrix-variate linear mixed model. Genetics 200:59–68

Gangurde SS, Wang H, Yaduru S, Pandey MK, Fountain JC, Chu Y, Isleib T, Holbrook CC, Xavier A, Culbreath AK (2020) Nested association mapping (NAM) based genetic dissection uncovers candidate genes for seed and pod weights in peanut (Arachis hypogaea). Plant Biotechnology Journal 18:1457–1471

Goudet J (2005) hierfstat, a package for r to compute and test hierarchical F-statistics. Molecular Ecology Notes 5:184–186

Guo M, Deng L, Gu J, Miao J, Yin J, Li Y, Fang Y, Huang B, Sun Z, Qi F, Dong W, Lu Z, Li S, Hu J, Zhang X, Ren L (2024) Genome-wide association study and development of molecular markers for yield and quality traits in peanut (Arachis hypogaea L.). BMC Plant Biol 24:244

Hill WG, Weir BS (1988) Variances and covariances of squared linkage disequilibria in finite populations. Theor Popul Biol 33:54–78

Huang L, Ren X, Wu B, Li X, Chen W, Zhou X, Chen Y, Pandey MK, Jiao Y, Luo H (2016) Development and deployment of a high-density linkage map identified quantitative trait loci for plant height in peanut (Arachis hypogaea L.). Scientific Reports 6:39478

Jahanzaib M, Nawaz N, Khurshid H, Jan SA, Arshad M, Hassan I (2019) Estimating genotype× environment interaction for groundnut seed yield across different ecological zones

Jiang Y, Xia B, Liang L, Li X, Xu S, Peng F, Wang R (2013) Molecular and analysis of a phenylalanine ammonia-lyase gene (LrPAL2) from Lycoris radiata. Molecular biology reports 40:2293–2300

Kafoutchoni KM, Agoyi EE, Agbahoungba S, Assogbadjo AE, Agbangla C (2021) Genetic diversity and population structure in a regional collection of Kersting’s groundnut (Macrotyloma geocarpum (Harms) Maréchal & Baudet). Genetic Resources and Crop Evolution 68:3285–3300

Lai K, Lorenc MT, Lee HC, Berkman PJ, Bayer PE, Visendi P, Ruperao P, Fitzgerald TL, Zander M, Chan CKK (2015) Identification and characterization of more than 4 million intervarietal SNP s across the group 7 chromosomes of bread wheat. Plant biotechnology journal 13:97–104

Li C, Guo L, Wang W, Miao P, Mu G, Chen CY, Meng C, Yang X (2024) Genome-Wide Identification, Characterization and Expression Profile of F-Box Protein Family Genes Shed Light on Lateral Branch Development in Cultivated Peanut (Arachis hypogaea L.). Horticulturae 10:255

Li J, Chen G-B, Rasheed A, Li D, Sonder K, Zavala Espinosa C, Wang J, Costich DE, Schnable PS, Hearne SJ, Li H (2019) Identifying loci with breeding potential across temperate and tropical adaptation via EigenGWAS and EnvGWAS. Molecular Ecology 28:3544–3560

Li J, Li D, Espinosa CZ, Pastor VT, Rasheed A, Rojas NP, Wang J, Varela AS, Carolina de Almeida Silva N, Schnable PS, Costich DE, Li H (2021) Genome-wide analyses reveal footprints of divergent selection and popping-related traits in CIMMYT’s maize inbred lines. J Exp Bot 72:1307–1320

Li L, Cui S, Dang P, Yang X, Wei X, Chen K, Liu L, Chen CY (2022) GWAS and bulked segregant analysis reveal the Loci controlling growth habit-related traits in cultivated Peanut (Arachis hypogaea L.). BMC genomics 23:403

Liu Y, Shao L, Zhou J, Li R, Pandey MK, Han Y, Cui F, Zhang J, Guo F, Chen J (2022) Genomic insights into the genetic signatures of selection and seed trait loci in cultivated peanut. Journal of advanced research 42:237–248

Lu Q, Huang L, Liu H, Garg V, Gangurde SS, Li H, Chitikineni A, Guo D, Pandey MK, Li S (2024) A genomic variation map provides insights into peanut diversity in China and associations with 28 agronomic traits. Nature genetics 56:530–540

Luo H, Xu Z, Li Z, Li X, Lv J, Ren X, Huang L, Zhou X, Chen Y, Yu J (2017) Development of SSR markers and identification of major quantitative trait loci controlling shelling percentage in cultivated peanut (Arachis hypogaea L.). Theoretical and Applied Genetics 130:1635–1648

Mallikarjuna N, Varshney RK (2014) Genetics, genomics and breeding of peanuts. CRC Press McAllister CH, Good AG (2015) Alanine aminotransferase variants conferring diverse NUE phenotypes in Arabidopsis thaliana. PLoS One 10:e0121830

Mo F, Zhang N, Qiu Y, Meng L, Cheng M, Liu J, Yao L, Lv R, Liu Y, Zhang Y (2021) Molecular characterization, gene evolution and expression analysis of the F-box gene family in tomato (Solanum lycopersicum). Genes 12:417

Oteng-Frimpong R, Karikari B, Sie EK, Kassim YB, Puozaa DK, Rasheed MA, Fonceka D, Okello DK, Balota M, Burow M (2023) Multi-locus genome-wide association studies reveal genomic regions and putative candidate genes associated with leaf spot diseases in African groundnut (Arachis hypogaea L.) germplasm. Frontiers in Plant Science 13:1076744

Otyama PI, Chamberlin K, Ozias-Akins P, Graham MA, Cannon EKS, Cannon SB, MacDonald GE, Anglin NL (2022) Genome-wide approaches delineate the additive, epistatic, and pleiotropic nature of variants controlling fatty acid composition in peanut (Arachis hypogaea L.). G3 (Bethesda) 12

Otyama PI, Kulkarni R, Chamberlin K, Ozias-Akins P, Chu Y, Lincoln LM, MacDonald GE, Anglin NL, Dash S, Bertioli DJ (2020) Genotypic characterization of the US peanut core collection. G3: Genes, Genomes, Genetics 10:4013–4026

Pandey MK, Upadhyaya HD, Rathore A, Vadez V, Sheshshayee M, Sriswathi M, Govil M, Kumar A, Gowda M, Sharma S (2014) Genomewide association studies for 50 agronomic traits in peanut using the ‘reference set’comprising 300 genotypes from 48 countries of the semi-arid tropics of the world. PLoS one 9:e105228

Pokrovac I, Pezer Ž (2022) Recent advances and current challenges in population genomics of structural variation in animals and plants. Frontiers in genetics 13:1060898

Poplin R, Ruano-Rubio V, DePristo MA, Fennell TJ, Carneiro MO, Van der Auwera GA, Kling DE, Gauthier LD, Levy-Moonshine A, Roazen D (2017) Scaling accurate genetic variant discovery to tens of thousands of samples. BioRxiv:201178

R Core Team (2016) R: a language and environment for statistical computing

Rafalski A, Ananiev E (2009) Genetic diversity, linkage disequilibrium and association mapping. Handbook of maize: genetics and genomics. Springer, pp 201–219

Raza A, Chen H, Zhang C, Zhuang Y, Sharif Y, Cai T, Yang Q, Soni P, Pandey MK, Varshney RK, Zhuang W (2024) Designing future peanut: the power of genomics-assisted breeding. Theoretical and Applied Genetics 137:66

Renaut S, Owens GL, Rieseberg LH (2014) Shared selective pressure and local genomic landscape lead to repeatable patterns of genomic divergence in sunflowers. Molecular Ecology 23:311–324

Sahu A, Rangari SK, Naik YD, Jyotish A, Pandey MK, Varshney RK, Thudi M, Punnuri SM (2025) Consensus genomic regions and key genes for biotic, abiotic and key nutritional traits identified using meta-QTL analysis in peanut. Frontiers in plant science 16:1539641

Sarlikioti V, de Visser PH, Buck-Sorlin G, Marcelis L (2011) How plant architecture affects light absorption and photosynthesis in tomato: towards an ideotype for plant architecture using a functional–structural plant model. Annals of Botany 108:1065–1073

Shah P, Pandey M, Nayak SN, Chen C, Bera S, Kole C, Puppala N (2023) Next-Generation Breeding for Nutritional Traits in Peanut. Compendium of Crop Genome Designing for Nutraceuticals. Springer, pp 403–417

Shaibu AS, Sneller C, Motagi BN, Chepkoech J, Chepngetich M, Miko ZL, Isa AM, Ajeigbe HA, Mohammed SG (2020) Genome-wide detection of SNP markers associated with four physiological traits in groundnut (Arachis hypogaea L.) mini core collection. Agronomy 10:192

Sharma R, Cockram J, Gardner KA, Russell J, Ramsay L, Thomas WT, O’Sullivan DM, Powell W, Mackay IJ (2021) Trends of genetic changes uncovered by Env-and Eigen-GWAS in wheat and barley. Theoretical and Applied Genetics:1–12

Sun Z, Qi F, Liu H, Qin L, Xu J, Shi L, Zhang Z, Miao L, Huang B, Dong W (2022) QTL mapping of quality traits in peanut using whole-genome resequencing. The crop journal 10:177–184

Upadhyaya HD, Ortiz R, Bramel PJ, Singh S (2003) Development of a groundnut core collection using taxonomical, geographical and morphological descriptors. Genetic Resources and Crop Evolution 50:139–148

Varshney RK, Pandey MK, Puppala N (2017) Future Prospects for Peanut Improvement. In: Varshney RK, Pandey MK, Puppala N (eds) The Peanut Genome. Springer International Publishing, Cham, pp 165–169

Venuprasad R, Aruna R, Nigam S (2011) Inheritance of traits associated with seed size in groundnut (Arachis hypogaea L.). Euphytica 181:169–177

Wang J, Yan C, Li Y, Li C, Zhao X, Yuan C, Sun Q, Shan S (2019) GWAS discovery of candidate genes for yield-related traits in peanut and support from earlier QTL mapping studies. Genes 10:803

Xie W, Wang G, Yuan M, Yao W, Lyu K, Zhao H, Yang M, Li P, Zhang X, Yuan J, Wang Q, Liu F, Dong H, Zhang L, Li X, Meng X, Zhang W, Xiong L, He Y, Wang S, Yu S, Xu C, Luo J, Li X, Xiao J, Lian X, Zhang Q (2015) Breeding signatures of rice improvement revealed by a genomic variation map from a large germplasm collection. Proceedings of the National Academy of Sciences 112:E5411–E5419

Zhang X, Zhang J, He X, Wang Y, Ma X, Yin D (2017) Genome-wide association study of major agronomic traits related to domestication in peanut. Frontiers in Plant Science 8:1611

Zhang X, Zhu L, Ren M, Xiang C, Tang X, Xia Y, Song D, Li F (2023) Genome-wide association studies revealed the genetic loci and candidate genes of pod-related traits in peanut (Arachis hypogaea L.). Agronomy 13:1863

Zheng X, Levine D, Shen J, Gogarten SM, Laurie C, Weir BS (2012) A high-performance computing toolset for relatedness and principal component analysis of SNP data. Bioinformatics 28:3326–3328

Zheng Z, Sun Z, Qi F, Fang Y, Lin K, Pavan S, Huang B, Dong W, Du P, Tian M (2024) Chloroplast and whole-genome sequencing shed light on the evolutionary history and phenotypic diversification of peanuts. Nature Genetics 56:1975–1984

Zhou X, Guo J, Pandey MK, Varshney RK, Huang L, Luo H, Liu N, Chen W, Lei Y, Liao B (2021) Dissection of the genetic basis of yield-related traits in the Chinese peanut mini-core collection through genome-wide association studies. Frontiers in plant science 12:637284

Zhou X, Stephens M (2012) Genome-wide efficient mixed-model analysis for association studies. Nature Genetics 44:821–824

Zhuang W, Chen H, Yang M, Wang J, Pandey MK, Zhang C, Chang W-C, Zhang L, Zhang X, Tang R (2019) The genome of cultivated peanut provides insight into legume karyotypes, polyploid evolution and crop domestication. Nature genetics 51:865–876

